# Proteomic analysis of the *Pseudomonas aeruginosa* iron starvation response reveals PrrF sRNA-dependent regulation of amino acid metabolism, iron-sulfur cluster biogenesis, motility, and zinc homeostasis

**DOI:** 10.1101/477984

**Authors:** Cassandra E. Nelson, Weiliang Huang, Luke K. Brewer, Angela T. Nguyen, Maureen A. Kane, Angela Wilks, Amanda G. Oglesby-Sherrouse

**Author notes:** Current Address: Silliker Food Science Center, 3600 Eagle Nest Drive, Crete, IL 60417. Equal contributions (Cassandra E. Nelson, Weiliang Huang).

## Abstract

Iron is a critical nutrient for most microbial pathogens, and the innate immune system exploits this requirement by sequestering iron and other metals through a process termed nutritional immunity. The opportunistic pathogen *Pseudomonas aeruginosa* provides a model system for understanding the microbial response to host iron depletion, as this organism exhibits a high requirement for iron as well as an exquisite ability to overcome iron deprivation during infection. Hallmarks of *P. aeruginosa’s* iron starvation response include the induction of multiple high affinity iron acquisition systems and an “iron sparing response” that is post-transcriptionally mediated by the PrrF small regulatory RNAs (sRNAs). Here, we used liquid chromatography-tandem mass spectrometry to conduct label-free proteomics of the *P. aeruginosa* iron starvation response, revealing several iron-regulated processes that have not been previously described. Iron starvation induced multiple proteins involved in branched chain and aromatic amino acid catabolism, providing the capacity for iron-independent entry of carbons into the TCA cycle. Proteins involved in sulfur assimilation and cysteine biosynthesis were reduced upon iron starvation, while proteins involved in iron-sulfur cluster biogenesis were paradoxically increased, highlighting the central role of iron in *P. aeruginosa* metabolism. Iron starvation also resulted in changes in the expression of several zinc-responsive proteins, as well as the first experimental evidence for increased levels of twitching motility proteins upon iron starvation. Subsequent proteomics analyses demonstrated that the PrrF sRNAs were required for iron regulation of many of these newly-identified proteins, and we identified PrrF complementarity with mRNAs encoding key iron-regulated proteins involved in amino acid metabolism, iron-sulfur biogenesis, and zinc homeostasis. Combined, these results provide the most comprehensive view of the *P. aeruginosa* iron starvation response to date and outline novel roles for the PrrF sRNAs in the *P. aeruginosa* iron sparing response and pathogenesis.

**AUTHOR SUMMARY:** Iron is central for the metabolism of almost all microbial pathogens, and as such this element is sequestered by the host innate immune system to restrict microbial growth. Defining the response of microbial pathogens to iron starvation is therefore critical for understanding how pathogens colonize and propagate within the host. The opportunistic pathogen *Pseudomonas aeruginosa*, which causes significant morbidity and mortality in compromised individuals, provides an excellent model for studying this response due to its high requirement for iron yet well-documented ability to overcome iron starvation. Here we used label-free proteomics to investigate the *P. aeruginosa* iron starvation response, revealing a broad landscape of metabolic and metal homeostasis changes that have not previously been described. We further provide evidence that many of these processes are regulated through the iron responsive PrrF small regulatory RNAs, which are integral to *P. aeruginosa* iron homeostasis and virulence. These results demonstrate the power of proteomics for defining stress responses of microbial pathogens, and they provide the most comprehensive analysis to date of the *P. aeruginosa* iron starvation response.

## INTRODUCTION

Iron is an essential micronutrient for nearly all forms of life, and presents a central paradigm for nutritional immunity, whereby the host sequesters iron from invading microbial pathogens. (1, 2). In turn, pathogens express a variety of high affinity iron acquisition systems to scavenge host iron (3, 4). In aerobic environments, iron poses the potential for toxicity through the production of reactive oxygen species (5). To balance the essentiality of iron with its potential for toxicity, bacteria must regulate the uptake, use, and storage of this nutrient in response to iron availability. In iron-replete conditions, iron uptake systems are repressed, while proteins involved in iron storage and oxidative stress protection are induced (6). Upon iron starvation, iron uptake systems are upregulated, while non-essential iron containing proteins are repressed in a strategy referred to as the iron-sparing response (7). Due to the central role of iron in numerous metabolic pathways, the iron sparing response is likely to elicit a substantial reorganization of bacterial metabolic networks. However, the full impact of iron starvation on bacterial metabolism, and how these changes impact pathogenesis, remain unclear.

As a pathogen with a substantial metabolic requirement for iron, *Pseudomonas aeruginosa* is an ideal model to elucidate the metabolic adaption to low iron starvation and the subsequent impact of this response on pathogenesis. *P. aeruginosa* is a Gram-negative opportunistic pathogen of significant concern for hospital-acquired infections, diabetic foot wound infections, and cystic fibrosis lung infections (8–11). To overcome iron limitation in the host, *P. aeruginosa* induces the expression of numerous exotoxins and proteases that cause tissue damage and may release host cell iron stores (12–14), as well as multiple high affinity iron acquisition systems to scavenge iron from host iron-sequestering proteins (15). In aerobic environments, iron can be acquired via the synthesis and secretion of two distinct siderophores, pyoverdine and pyochelin, which scavenge oxidized ferric iron (Fe^3+^) from host proteins such as transferrin and lactoferrin (16). In anaerobic environments, reduced ferrous iron (Fe^2+^) is acquired through the inner membrane associated Feo transport system (17). In addition to these labile iron transport systems, *P. aeruginosa* utilizes two non-redundant heme uptake systems, Has (heme assimilation system) and Phu (Pseudomonas heme uptake), to acquire host heme as an iron source (18). Studies have demonstrated a role for each of these systems in different infection models (19–22), highlighting iron as a central mediator of *P. aeruginosa* pathogenesis.

Iron starvation also induces expression of the PrrF small regulatory RNAs (sRNAs), which we have shown are required for acute murine lung infection (23). The PrrF sRNAs post-transcriptionally repress the expression of multiple iron-containing proteins, presumably by pairing with, destabilizing, and reducing translation of the encoding mRNAs (24, 25). Included in the PrrF regulon are iron-and heme-cofactored components of the tricarboxylic acid (TCA) cycle and oxidative phosphorylation pathways, which are central components of *P. aeruginosa* metabolism (24, 25). *P. aeruginosa* is able to grow well in low iron environments in spite of this downregulation, yet the strategies employed to compensate for the loss of these systems have yet to be identified (23). We postulate that this gap in knowledge is due in part to the previous reliance on RNA-centric approaches, such as microarray analyses, which are only able to identify transcriptional and some post-transcriptional regulatory effects of iron limitation. In contrast, proteomics can reveal the full scope of transcriptional, post-transcriptional, translational, and post-translational regulatory changes due to iron depletion, including those mediated by the PrrF sRNAs, providing a more comprehensive analysis of iron-dependent regulation.

Herein we used liquid chromatography-tandem mass spectrometry (LC-MS/MS) to quantify changes in the *P. aeruginosa* proteome in iron-depleted conditions. In addition to the expected decreases in proteins in the TCA cycle and oxidative phosphorylation, iron starvation induced a compensatory increase in proteins for the catabolism of branched chain and aromatic amino acids. Iron depletion also reduced the levels of proteins for sulfur assimilation and increased levels of proteins for iron-sulfur (Fe-S) cluster biogenesis. We further identified novel iron regulation of several zinc-responsive proteins, and we showed that iron depletion increases levels of proteins for twitching motility. Subsequent proteomic analysis of the ∆*prrF* mutant showed that iron regulation of several of these pathways is partially dependent on the PrrF sRNAs. Moreover, we identified PrrF complementarity with mRNAs encoding several novel iron-regulated proteins, suggesting direct post-transcriptional regulation of these pathways. These results provide the most comprehensive analysis of the *P. aeruginosa* iron starvation response to date, revealing novel strategies likely used by this pathogen to survive in the host and initiate infection.

## RESULTS

### Proteomics reveals broad changes in metabolic proteins upon iron starvation

To analyze protein expression changes in *P. aeruginosa* under high and low iron conditions, we performed quantitative label-free proteomics on reference strain PAO1 cultures grown in chelex-treated dialyzed tryptic soy broth (DTSB) supplemented with or without 100µM FeCl3. These growth conditions have been used for numerous *P. aeruginosa* iron regulatory studies, including previous GeneChip analyses of the iron starvation response (26) and PrrF regulation (24, 25). Moreover, DTSB is rich in amino acids, which have been shown to be abundant in disease states such as the cystic fibrosis (CF) lung and to promote the production of numerous iron-regulated virulence factors (27–29). Protein samples were purified preliminarily by size exclusion ultrafiltration, cleaved by trypsin, and subjected to differential expression analysis by nanoLC-ion mobility linked parallel MS, termed ultradefinition MS^e^ (UDMS^e^). UDMS^e^ is a data-independent acquisition method utilizing tandem mass spectrometry with traveling wave ion mobility that uses ion mobility drift time-specific collision energy profiles to enhance precursor fragmentation and depth of coverage (30). Subsequent informatics were used to identify proteins that were significantly (*p* < 0.05) induced or repressed at least two-fold (equivalent to 1 log_2_ fold change, or LFC) upon iron starvation. The complete results of this analysis are provided in the supplementary materials (**Dataset S1**).

In total, we identified 309 proteins as induced by iron starvation, while 250 proteins were repressed upon iron starvation. Of these proteins, 240 of the induced and 226 of the repressed proteins upon iron starvation have not been reported as iron regulated in previously published studies (24–26, 31). Pathway analysis was performed on the KEGG database to determine what cellular activities and metabolic pathways were likely affected by changes in the iron-induced and iron-repressed proteins. As expected, these results demonstrated that iron starvation repressed proteins in numerous metabolic pathways. In contrast, proteins involved in iron acquisition, ketone body metabolism, and Type IV pilus-dependent motility were significantly increased under low iron conditions (**Table 1**). While some of these pathways were previously known to be iron-responsive, iron regulation of proteins in ketone body metabolism and twitching motility has not been described. A closer examination of this dataset as described below revealed several potential mechanisms for adapting to iron starvation, including changes in proteins for amino acid metabolism, Fe-S cluster biogenesis, and zinc homeostasis.

**Table 1.** Iron regulated pathways of PAO1 identified through pathway analysis

| Pathway | p-value |
| --- | --- |
| **Downregulated in low iron** |  |
| Oxidative phosphorylation | $3 \times 10^{-14}$ |
| Microbial metabolism in diverse environments | $1 \times 10^{-4}$ |
| Tricarboxylic Acid Cycle | $3 \times 10^{-3}$ |
| Aminobenzoate degradation | $3 \times 10^{-3}$ |
| Metabolic pathways | $3 \times 10^{-3}$ |
| Polycyclic aromatic hydrocarbon degradation | $4 \times 10^{-2}$ |
| **Upregulated in low iron** |  |
| Heme assimilation and utilization | $4 \times 10^{-18}$ |
| Pyoverdine biosynthesis | $1 \times 10^{-14}$ |
| Pyochelin biosynthesis | $2 \times 10^{-4}$ |
| Ketone bodies synthesis and degradation | $4 \times 10^{-2}$ |
| Type IV pilus-dependent motility | $5 \times 10^{-2}$ |

### Proteomics verifies the classical *P. aeruginosa* iron starvation response

The *P. aeruginosa* transcriptional response to iron starvation has been well characterized (20, 24-26, 32). Expected changes include the upregulation of several iron uptake systems, including those for siderophore, heme, and ferrous iron uptake systems. In agreement with these previous studies, our results show that numerous proteins involved in siderophore-mediated iron uptake via pyoverdine and pyochelin were upregulated at least two-fold under low iron conditions (**Supplementary Materials, Table S1**). Key components of the heme acquisitions systems were also induced in low iron, including proteins in the heme acquisition system (HasR, HasA, and HasI), the heme outer membrane heme receptor PhuR, and the iron-regulated heme oxygenase HemO (**Supplementary Materials, Table S1**). Also in agreement with earlier transcriptional studies, we found that several iron-dependent proteins were down-regulated under iron starvation conditions, including the iron-cofactored superoxide dismutase SodB, the heme-cofactored catalase KatA, the iron storage protein bacterioferritin B, BfrB, and a putative bacterioferritin encoded by PA4880 (**Supplementary Materials, Table S1**). The anthranilate degradative enzymes AntABC, which utilize an iron cofactor, were also downregulated upon iron starvation (24, 25), while the iron-activated AntR transcriptional activator of the *antABC* genes (24) was not detected in this study.

While many of the previously-identified transcriptional responses to iron starvation were evident in this dataset, we also noted some discrepancies between protein level changes in our study and previously reported changes in RNA levels. Specifically, some iron-regulated integral membrane proteins, such as FpvJ, FpvE, PhuU, HasE, and HasD, were not detected in our analysis, possibly due to either the insolubility or low abundance of these proteins. Also as expected, extracellular proteins such as exotoxin A and the secreted hemophore HasA were not detected, as the secreted proteome was not analyzed in the current study. We also noted that some iron-regulated genes encoding cytoplasmic proteins were detected but did not exhibit iron regulation at the protein level. This included the HasS antisigma factor, which is predicted to regulate the Has heme assimilation system (33), and the AprA protease, which functions to modulate the host immune response (34) (**Supplementary Materials, Table S1**). One explanation for this discrepancy is that these proteins are subject to additional translational or post-translational regulatory mechanisms that mask the previously-observed transcriptional iron regulatory responses.

### *P. aeruginosa* induces multiple iron-sparing metabolic pathways during iron starvation

Previous microarray studies demonstrated reduced expression of multiple iron-containing TCA cycle enzymes under iron limiting conditions (24–26). In agreement with these studies, we observed a reduction in the TCA cycle enzymes citrate synthase (GltA), aconitase (AcnA and AcnB), and succinate dehydrogenase (SdhCDAB) (**Fig. 1A, and Supplementary Materials, Fig S1A**). However, it remained unclear how iron starved cells metabolically compensate for the reduced expression of these enzymes. One possibility was that *P. aeruginosa* shifts its metabolism to utilize the glyoxylate shunt, which was recently observed in iron depleted M9 medium (35). While we observed a significant upregulation of the isocitrate lyase AceA, we also observed a significant downregulation of the malate synthase GlcB under low iron conditions (**Supplementary Materials, Table S1**). Thus, it does not appear that the glyoxylate shunt is the primary means of metabolism under iron-starved conditions in an amino acid rich medium (**Fig 1A, and Supplementary Materials, Fig S1A**).

**Figure 1.**
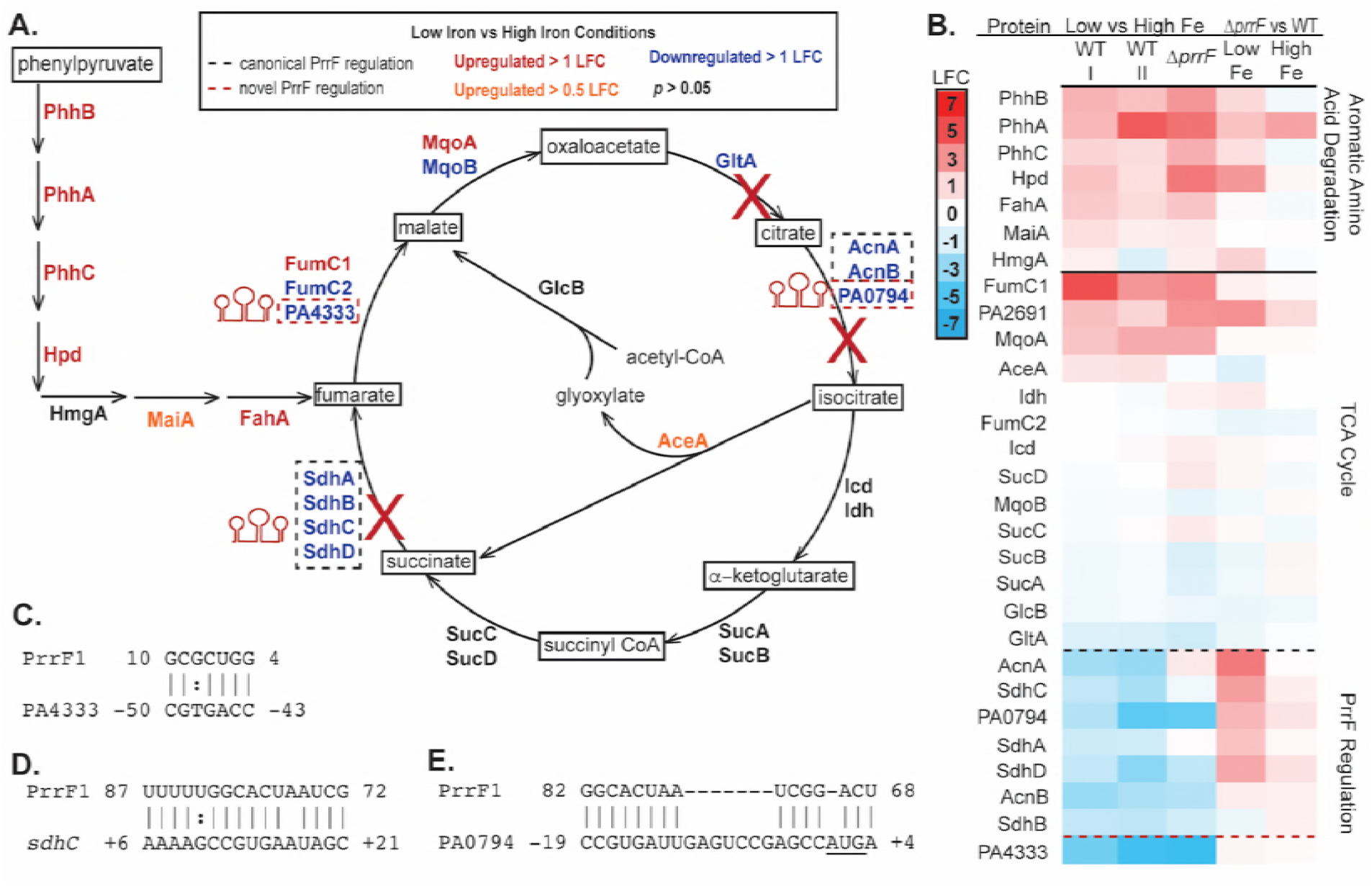
Proteins for aromatic amino acid degradation are increased in response to low iron conditions. **A)** Under iron-replete conditions, *P. aeruginosa* uses the TCA cycle to generate energy. Under low iron conditions, iron-containing proteins indicated by blue text are repressed, causing breaks in the TCA cycle, shown with red X’s. Proteomics revealed enzymes for aromatic amino acid catabolism are upregulated under low iron conditions, indicated by red or orange text, providing the capacity to produce the TCA cycle intermediate fumarate. A subsequent proteomics experiment was performed comparing expression under high and low conditions of PAO1 and ∆*prrF*, confirming previously identified PrrF regulation of these proteins. Detailed annotation of the metabolic intermediates is shown in the Supplementary Materials Figure S1. **B)** Heatmap showing changes in protein expression of aromatic amino acid degradation enzymes and TCA cycle enzymes from the first proteomics experiment analyzing wild type PAO1 (I) and subsequent experiment including wild type PAO1 and ∆*prrF* (II). Previously identified PrrF regulation is indicated with a black dashed line, and novel PrrF regulation is indicated with a red dashed line. **C-E)** Novel PrrF complementarity with the PA4333 (**C**), *sdhC* (**D**), and PA0794 (**E**) mRNAs was identified using CopraRNA (59).

Our data instead indicate that *P. aeruginosa* shifts to amino acid catabolism under low iron conditions. The enzymes that comprise the aromatic amino acid degradation pathway, which converts phenylpyruvate, phenylalanine, and tyrosine into fumarate, were almost all significantly upregulated above our two-fold threshold (**Fig 1A, and Supplementary Materials, Fig S1A**). Notably, the entry of these metabolites into the TCA cycle takes advantage of the iron-independent paralog of fumarate hydratase (FumC1) (36) and probable iron-independent paralog of malate:quinone oxidoreductase (MqoA), both of which were upregulated under low iron conditions (**Fig. 1A, and Supplementary Materials, Fig S1A**). Our pathway analysis also indicated proteins involved in ketone body metabolism, a process that degrades fatty acids and branched chain amino acids to allow fasting in higher organisms, were induced upon iron limitation. Ketone body metabolism has not been reported in prokaryotes, but we did observe increased levels of proteins for ketogenic amino acid metabolism. Specifically, several enzymes for leucine catabolism (Ldh, BkdA1, BkdB, LiuD, and LiuE) were significantly upregulated at least 1.5-fold (0.5 LFC) in low iron conditions (**Fig. 2A, and Supplementary Materials, Fig S2A**). Conversely, several proteins involved branched chain amino acid biosynthesis (IlvE, IlvD and LeuC) were down-regulated in low iron conditions (**Fig. 2B, and Supplementary Materials, Fig S2**). Combined, these data suggest *P. aeruginosa* shifts to amino acid catabolism upon iron starvation to support its metabolism.

**Figure 2.**
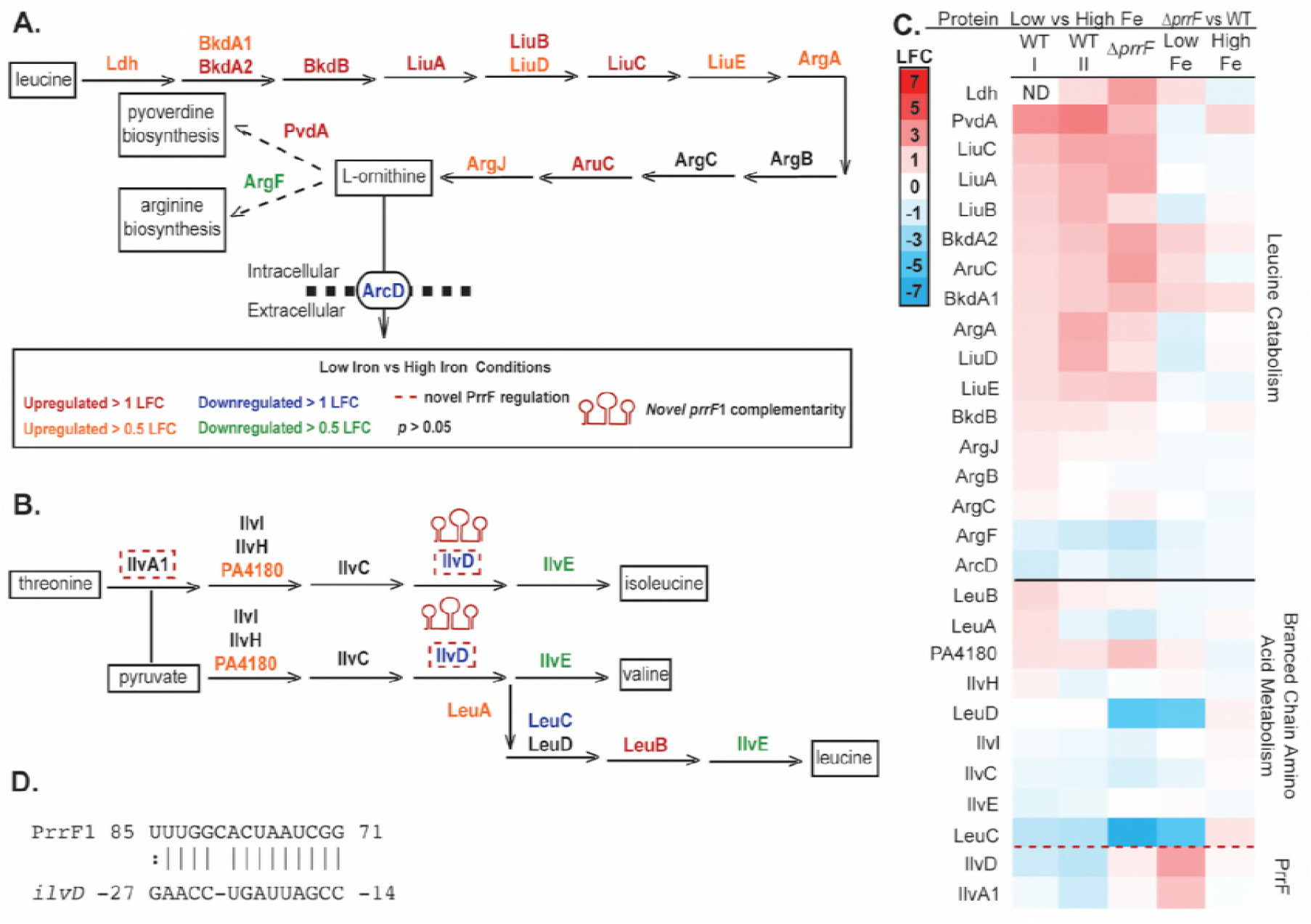
Proteins for branched chain amino acid metabolism and catabolism are regulated by iron. **A)** Proteomics revealed enzymes involved in leucine catabolism to L-ornithine are upregulated in low iron conditions, indicated by red and orange text. Ornithine may be spared for production of pyoverdine through the downregulation of the ornithine/arginine antiporter ArcD and ArgF, the first enzyme in arginine biosynthesis, indicated by blue and green text. **B)** The biosynthesis of the branched chain amino acids, leucine, threonine, and valine is seemingly reduced through the downregulation of multiple biosynthesis proteins in response to low iron. Detailed annotation of the metabolic intermediates for panels **A-B** is shown in the Supplementary Materials Figure S2. **C)** Heatmap showing changes in protein expression of branched chain amino acid biosynthesis and leucine degradation from the first proteomics experiment including only wild type PAO1 (I) and subsequent experiment including wild type PAO1 and ∆*prrF* (II). Novel PrrF regulation of IlvD and IlvA1 was identified, as indicated by a red dashed line. **D)** PrrF complementarity with the *ilvD* mRNA was identified using CopraRNA (59).

We also observed modest but significant increases in proteins that synthesize L-ornithine from glutamate and acetyl-CoA (ArgA, ArgB, ArgC, and AruC) (28) (**Fig 2A, and Supplementary Materials, Fig S2A**). The idea that the cell is metabolizing glutamate to form L-ornithine is attractive, as the DTSB medium used for this study is supplemented with an excess of mono-sodium glutamate as a nitrogen source, and ornithine is needed for pyoverdine production (22). Interestingly, the ArcD L-ornithine/arginine antiporter, and ArgF, which incorporates ornithine into the arginine biosynthesis pathway, were both significantly downregulated in iron-depleted conditions (**Fig 2A, and Supplementary Materials, Fig S2A**). This could be part of an ornithine sparing response for pyoverdine biosynthesis, further supported by the upregulation of the first pyoverdine biosynthetic enzyme PvdA (**Supplementary Materials, Table S1**). This possible metabolite sparing strategy to promote pyoverdine synthesis is similar to the previously-described down-regulation of the AntABC anthranilate degradation enzymes to promote 2-alky-4(1*H*)-quinolones (AQs) under low iron conditions (24, 37).

As previously reported, we also observed decreased levels of several iron-dependent proteins involved in oxidative phosphorylation in low iron conditions (**Supplementary Materials, Table S1**) (24–26). In contrast, levels of the lower affinity cytochrome ubiquinol oxidase CyoA, part of the CyoABCDE complex that is less reliant on iron (38), were increased in low iron (**Supplementary Materials, Table S1**). CyoBCDE were not detected, possibly due to their insolubility as integral membrane proteins. Low iron also resulted in a 1.7 LFC of PA2691 (**Supplementary Dataset S1**), which is a predicted type II NADH:quinone oxidoreductase (NDH-2). NDH-2 proteins utilize flavin cofactors instead of iron to catalyze the oxidation of NADH and reduce quinones (39). Thus, PA2691 may function in place of the *nuoA-N* encoded complex I to oxidize NADH under low iron conditions. These data outline a strategy wherein *P. aeruginosa* shifts production of enzymes for oxidative phosphorylation pathways to support iron-sparing respiratory metabolism.

### Iron regulates proteins for sulfur acquisition and metabolism

Sulfur is a key component of Fe-S clusters, which are incorporated into many metabolic enzymes. This and other studies have shown that numerous Fe-S containing proteins are downregulated during iron-depleted growth (24, 25). It is therefore possible that *P. aeruginosa* alters pathways for the uptake and metabolism of sulfur in response to changing iron availability. In agreement with this hypothesis, proteins with demonstrated and putative roles in alkanesulfonate uptake (SsuA, PA2594, and PA2595), as well as the alkanesulfonate monooxygenases PA2600 and SsuD, were reduced in low iron conditions (**Fig. 3A, and Supplementary Materials, Fig S3A-B**). Moreover, iron limitation resulted in decreased levels of several proteins involved in cysteine biosynthesis from sulfate, including CysD, CysN, CysI, CysH, and PA2600 (**Fig. 3A, and Supplementary Materials, Fig S3A-B**). While the CysK and CysM enzymes that catalyze the final steps of cysteine biosynthesis were not affected by iron depletion in our study, decreases in proteins that mediate the earlier steps in this pathway strongly suggest that *P. aeruginosa* reduces sulfur assimilation and cysteine biosynthesis pathways under low iron conditions.

**Figure 3.**
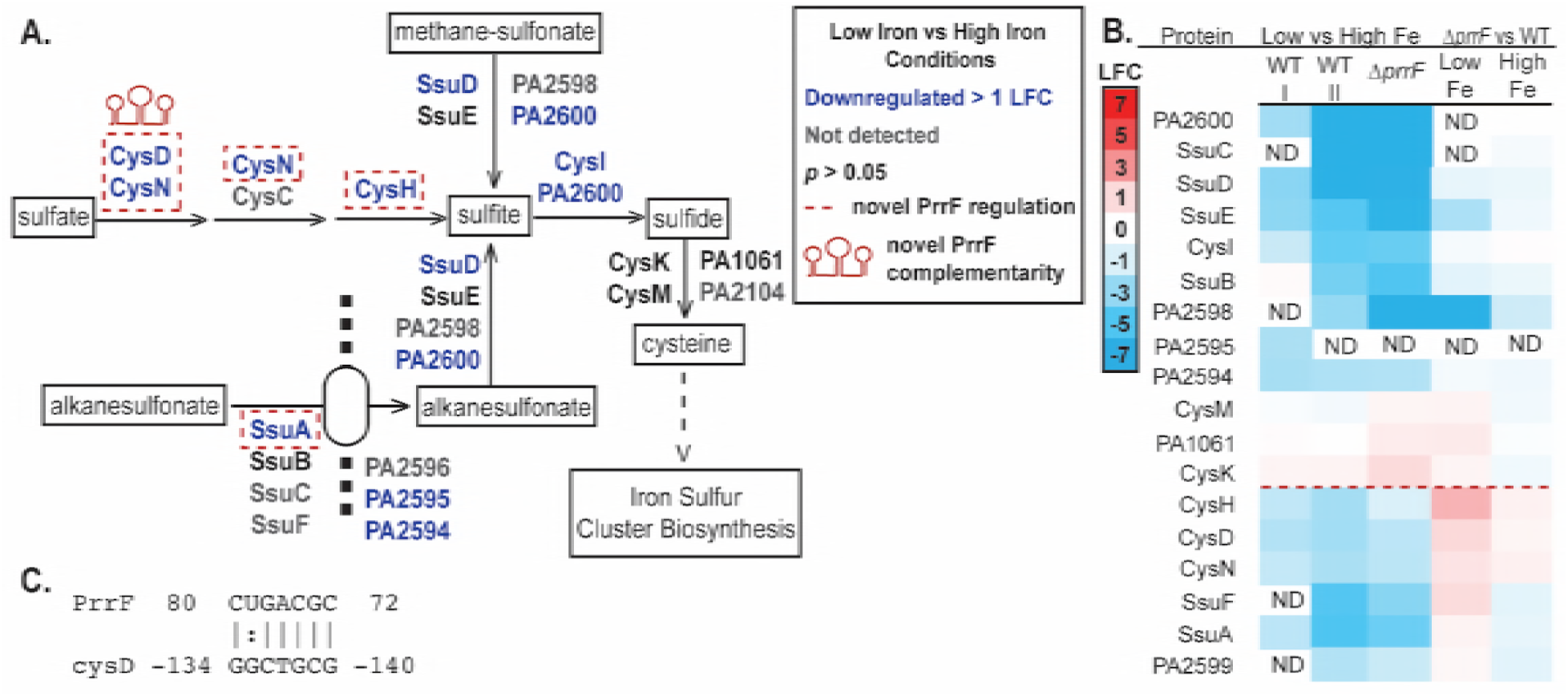
Proteomics reveals iron regulation of proteins for sulfur assimilation and cysteine biosynthesis. **A)** Schematic of cysteine biosynthesis. Sulfide for cysteine biosynthesis can be generated from imported extracellular alkanesulfonate, intracellular sulfate, or intracellular methane-sulfonate. Key proteins involved in this process were identified as down-regulated during iron limitation (blue text), indicating a decrease in cysteine production. **B)** Heatmap showing changes in protein expression of cysteine biosynthesis enzymes from the first proteomics experiment with wild type PAO1 (I) and subsequent experiment with wild type PAO1 and ∆*prrF* (II). Novel PrrF regulation was identified for six of the enzymes and separated with a red dashed line. **C)** Complementarity between the PrrF sRNA and the UTR of the *cysD* mRNA was identified by CopraRNA (59).

Interestingly, almost all of the known proteins involved with Fe-S cluster biosynthesis, including IcsR, HscAB, IcsX, and the potential iron donor CyaY (40), were significantly upregulated in iron-depleted conditions (**Fig. 4**). In contrast, IcsU, which is encoded on the same operon as other Fe-S synthesis proteins, was significantly downregulated in iron-depleted conditions. Differential impacts of iron on proteins encoded by the *Isc-hsc-fdx* operon may be due to either dis-coordinate post-transcriptional regulatory activities or differences in protein half-life. We also identified a partial Suf-like locus encoding a SufS-like desulfurase, PA3668, and a SufE-like sulfur transport protein, PA3667, both of which were significantly upregulated in low iron conditions (**Fig. 4**). The upregulation of the SufE-like protein (PA3667) may be able to compensate for the reduced levels of IscU in low iron conditions (**Fig. 4B**). Overall, these results indicate that *P. aeruginosa* upregulates Fe-S cluster biosynthesis proteins in low iron conditions, despite the global downregulation of Fe-S cluster containing proteins. This response to iron starvation has previously been reported for the *suf* locus in *Escherichia coli* (41), and is thought to be necessary to maintain Fe-S cluster biogenesis for essential proteins. Thus, these results suggest a central requirement for Fe-S cluster biogenesis in *P. aeruginosa* metabolism even under conditions of iron starvation.

**Figure 4.**
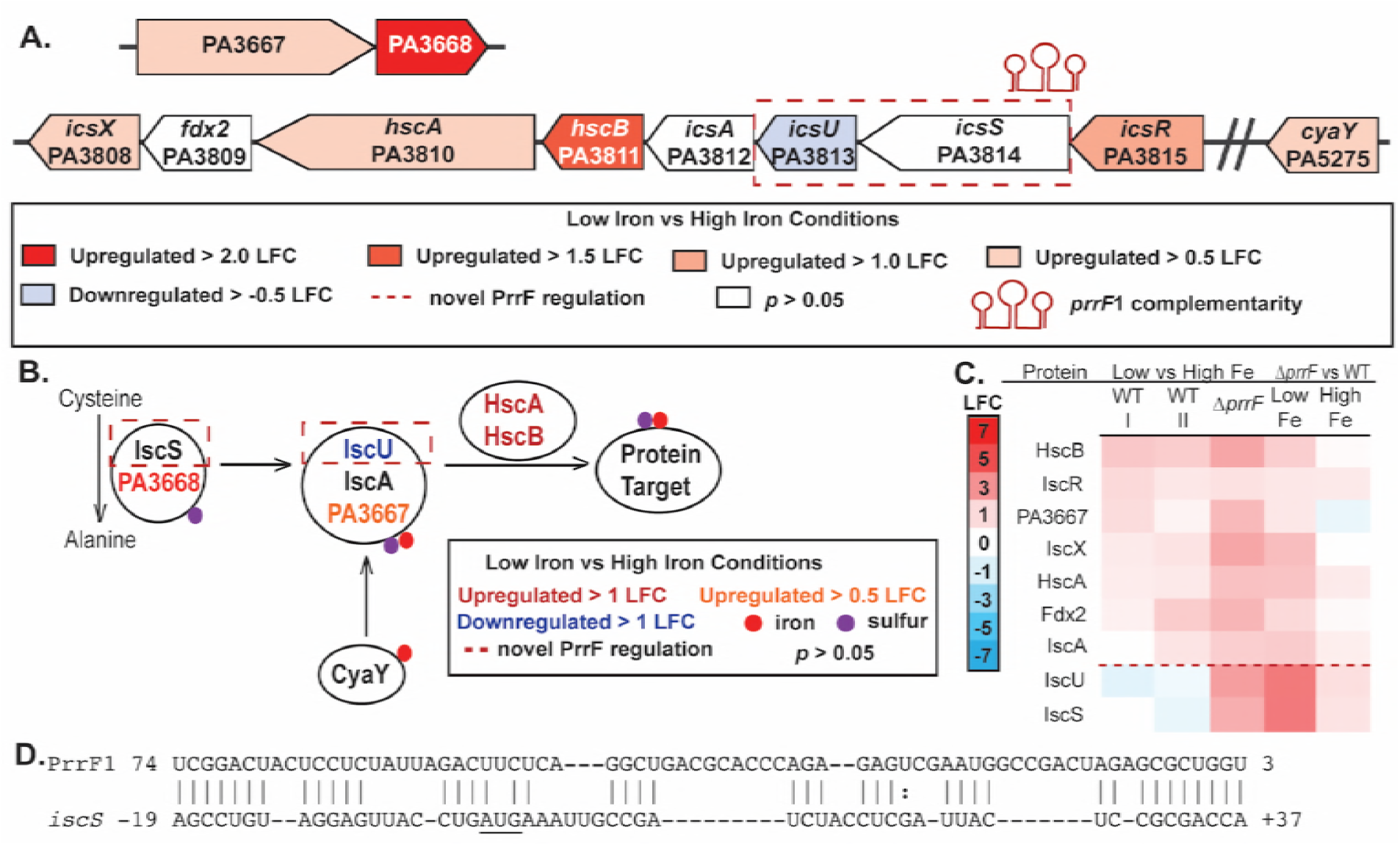
Proteomics reveals complex regulation of Fe-S cluster biogenesis. **A)** The *isc* gene cluster is transcribed as an operon from *icsR* to *icsX* (68), yet our results show only IcsU is specifically downregulated under low iron conditions (indicated by blue shading). An additional Suf-like locus was identified and found to be upregulated under low iron conditions (indicated by orange shading). A homolog to the eukaryotic frataxin, *cyaY,* is encoded separately on the chromosome. **B)** Model of Fe-S generation. Cysteine is defulonated by IscS or PA3668 to form alanine. The sulfur is then passed to the sulfur transfer proteins IscU, IscA, or PA3667. An iron atom is donated, possibly by CyaY, to form the Fe-S cluster. The chaperones HscA and HscB transfer the Fe-S cluster to the protein target. Steps subject to PrrF-mediated iron regulation are indicated by a red dashed box. **C)** Heatmap showing log-fold changes (LFC) in the levels of Fe-S biogenesis proteins from the first proteomics experiment with wild type PAO1 (I) and subsequent experiment including wild type PAO1 and ∆*prrF* (II). Novel PrrF regulation of IscS and IscU was identified and denoted with a red dashed line. **D)** Complementarity between PrrF and the *icsS* mRNA was identified by CopraRNA (59).

### Proteins for phenazine biosynthesis are repressed in iron-poor conditions

Phenazines are redox active secondary metabolites that contribute to biofilm formation, act as extracellular electron acceptors, and exhibit antimicrobial properties (42–45). Biosynthesis of phenazine-1-carboyxilic acid (PCA) from chorismate is performed by enzymes encoded by two almost identical operons, *phzA1-phzG1* (*phz1)* and *phzA2-G2* (*phz2)* (46) (**Fig 5A**). The two redundant operons have been shown to be regulated independently and have different roles in pathogenicity, with the *phz2* operon required and sufficient for pathogenesis (48). PCA can be further modified by PhzH to produce phenazine-1-carboxamide (PCN), by PhzS to produce 1-hydroxyphenazine (1-OH-PHZ), or by PhzM to produce 5-methylphenazine-carboxylic acid (5-Me-PCA), from which pyocanin (PYO) is produced by PhzS (**Fig. 5A**). Many but not all of these biosynthetic proteins, as well as the MexGHI-OmpD multi-drug efflux pump that is required for secreting phenazines into the environment, were significantly reduced in low iron conditions (**Fig. 5A**) (47). The amino acid sequences of the enzymes encoded by the two operons are 100% identical from PhzC-PhzG, so only PhzA1/A2 and PhzB1/B2 could be differentiated by proteomic analysis. Though unique peptides were identified to distinguish the PhzA1 and PhzA2 proteins, the extracted-ion chromatogram (XIC) quality of these unique peptides was not sufficient for differential quantification. Therefore we differentiated the expression of the two operons using PhzB expression: PhzB2 was significantly repressed under low iron conditions, while PhzB1 was not affected by iron (**Fig. 5C**). Based on these data, we hypothesize that iron specifically regulates proteins encoded by the *phz2* operon, though further studies will be necessary to thoroughly test this hypothesis.

**Figure 5.**
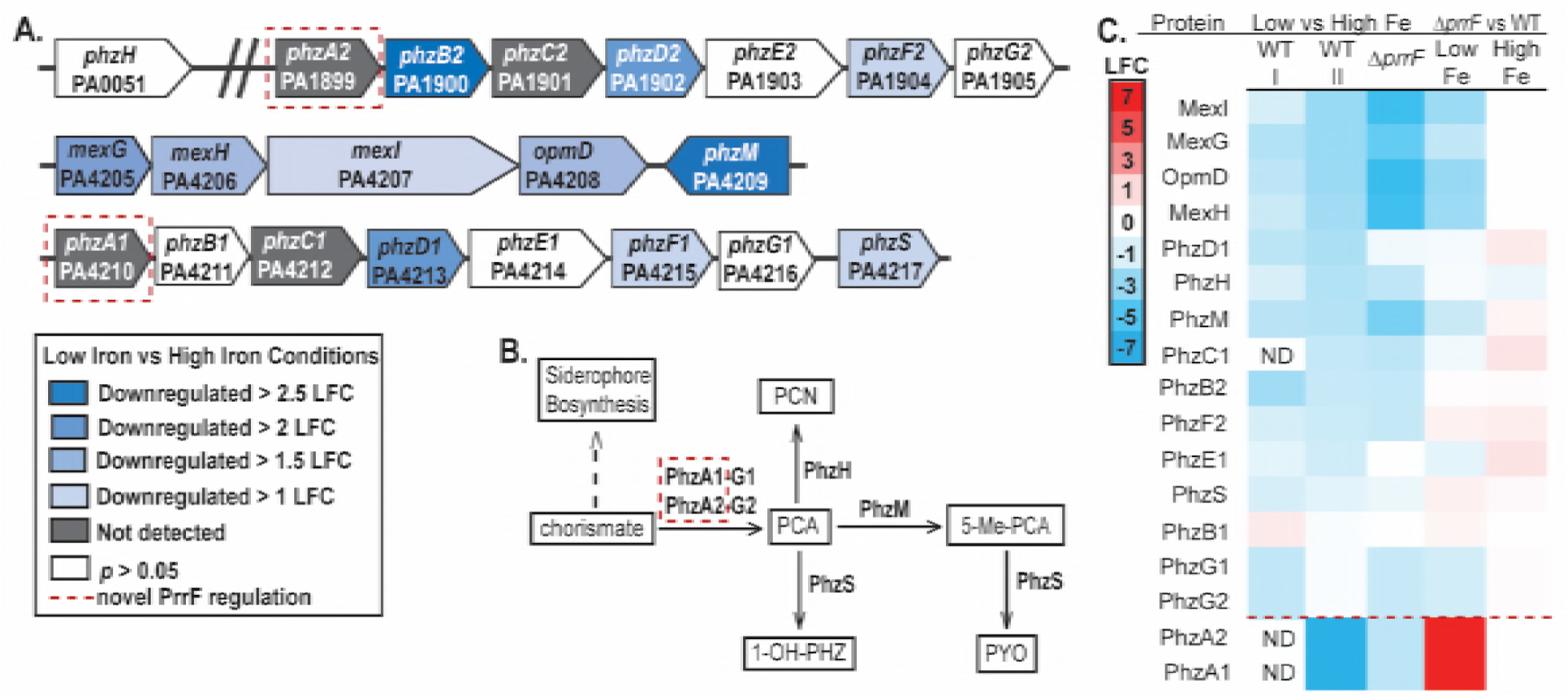
Proteins for phenazine biosynthesis are downregulated under low iron conditions. **A)** Organization of the phenazine biosynthetic enzymes and efflux pump genetic organization. Protein expression fold change under low iron conditions compared to high iron conditions is represented by color. **B)** Phenazine biosynthesis pathway. Phenazine carboxylic acid (PCA) is generated from chorismate by the enzymes encoded in the almost identical *phzA1-G1* and *phzA2-G2* operons. PCA can be further modified by PhzH, PhzM and PhzS to form phenazine-1-carboxyamide (PCN), 5-methylphenazine-1-carboxylic acid (5-Me-PCA), and 1-hydroxyphenazine (1-OH-PHZ) respectively. 5-Me-PCA can be further modified to form pyocyanin (PYO). PrrF-dependent iron regulation of the PhzA proteins was identified in the current study and is indicated by a red dashed box. **C)** Heatmap showing log-fold changes (LFC) in phenazine biosynthesis and transport protein expression from the first proteomics experiment with wild type PAO1 (I) and subsequent experiment including wild type PAO1 and ∆*prrF* (II). Novel PrrF regulation was identified for PhzA1 and PhzA2 and denoted with a red dashed line.

### Structural and regulatory proteins for twitching motility are increased under low iron conditions

Previous work demonstrates that *P. aeruginosa* limits motility in high iron conditions to promote biofilm formation, while iron starvation induces twitching motility (49, 50). However, microarray studies have not identified iron regulation of the genes that encode the twitching motility apparatus (26). Here we show that many of the proteins comprising the pilus required for twitching motility are significantly increased upon iron starvation. This includes the PilM and PilN proteins in the alignment complex, the major pilus subunit PilA, and almost all of the minor pillin subunits (**Fig.6A**). Several of the proteins involved in the twitching-specific chemosensory system, including PilG, PilH, and ChpC, are also upregulated in iron-depleted conditions (**Fig.6A**). To our knowledge, this is the first demonstration of iron-regulated levels of the proteins involved in twitching motility.

The induction of twitching motility proteins in this experiment was particularly interesting as this experiment was performed with shaking cultures, and pili-mediated twitching motility occurs during static growth on solid or semi-solid surfaces. To confirm that iron would have a similar effect on twitching motility on solid DTSB medium, twitching motility assays were performed in DTSB agar plates with and without 100uM iron supplementation. In agreement with previous studies, the twitching diameter for wild type PAO1 was significantly larger under low iron conditions than high iron conditions (**Fig. 6B and Supplementary Materials, Fig. S3**). Combined, these data demonstrate that increased twitching in low iron conditions is likely due to increased levels of many components of the Type IVa pili machinery.

**Figure 6.**
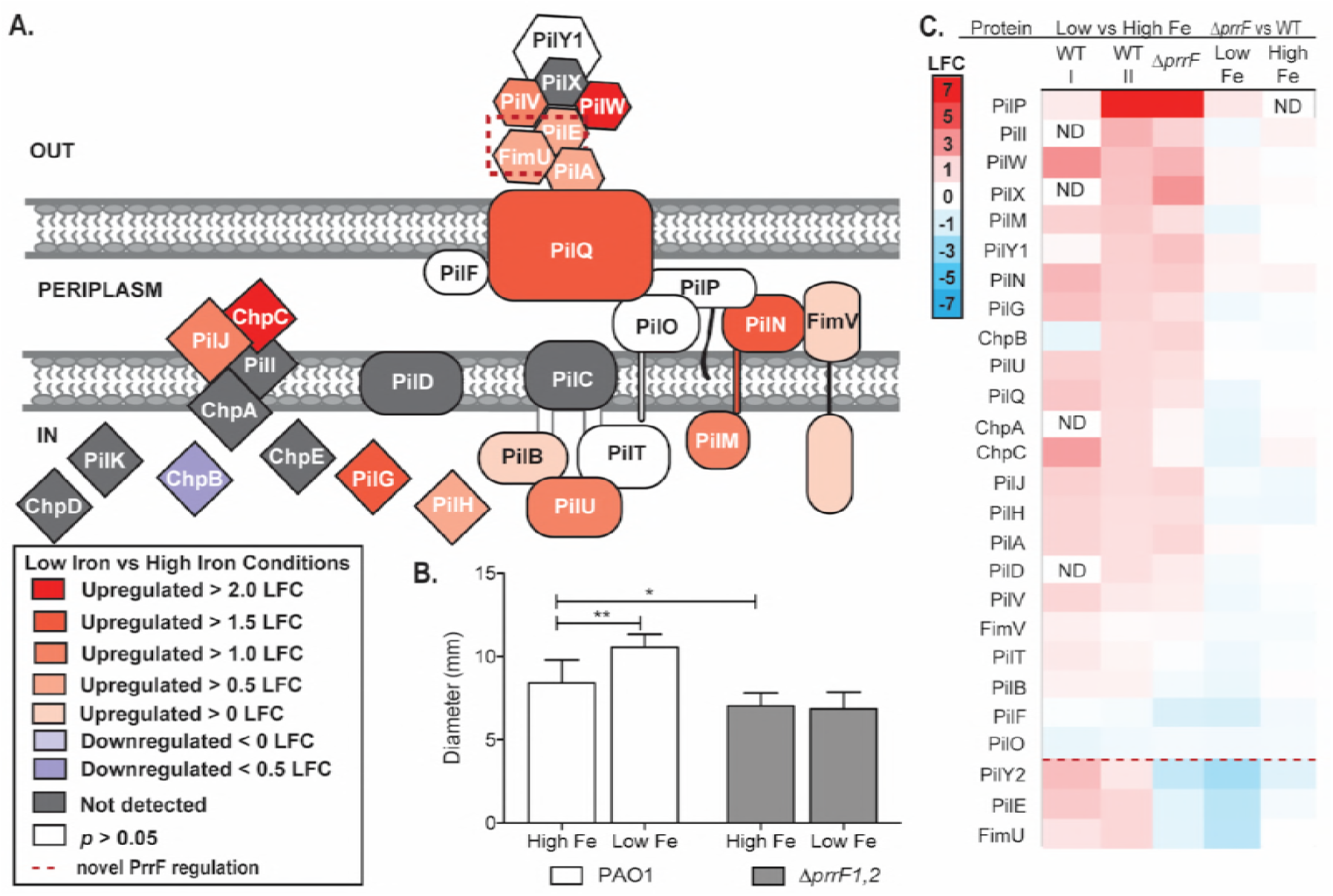
Mechanism for iron regulation of twitching motility is revealed by proteomics. **A)** Proteomics identified iron-regulated expression of the major pillin protein PilA, and the minor pillin proteins PilV, PilW, PilW, and FimU. Iron regulation was also identified for most of the assembly subcomplex, which is comprised of PilF, PilQ, PilB, PilC, PilD, PilT, and PilU, and the alignment subcomplex, comprised of FimV, PilP, PilO, PillN, and PilM. The pilus specific chemotaxis system (Pil-Chp), comprised of PilHIJK and ChpABCDE, regulates twitching motility chemotaxis. Upregulation under low iron conditions was identified for PilJ, PilG, PilH, and ChpC, indicated by orange shading of the proteins, while the expression of ChpB was downregulated, indicated by blue shading of the protein. **B)** Twitching motility assays were performed for PAO1 and ∆*prrF* and quantified after staining with 1% crystal violet. Error bars represent the standard deviation of 10 biological replicates in technical duplicate. Significance was determined using a Students two tailed T-test *: *p* = 0.01 **: *p* < 0.005. **C)** Heatmap showing log-fold changes (LFC) in the levels of twitching motility proteins from the first proteomics experiment with wild type PAO1 (I) and subsequent experiment including wild type PAO1 and ∆*prrF* (II).

### Proteomics reveals regulatory crosstalk between iron and zinc

As discussed above, iron starvation in *P. aeruginosa* results in the induction of iron-independent paralogs of certain metabolic enzymes (*e.g.* FumC and Mqo-mediated reactions in **Fig. 1A**). In some cases, these enzymes rely on other transition metals, such as zinc and manganese, to support their structure or activity. Thus, one possible strategy to compensate for iron starvation may be to induce pathways for the uptake of other transition metal ions. In support of this idea, our data demonstrate that iron limitation results in increased levels of proteins encoded by the *cnt* operon, which mediate the synthesis, secretion, and uptake of a novel opine metallophore involved in zinc uptake (**Fig 7A-B**). Interestingly, CntL, the nicotianamine synthase (51), was not statistically increased by iron depletion, and CntI, the predicted cytoplasmic membrane exporter of pseudopaline (52), was not detected. Our finding that CntL protein levels are not affected by iron depletion is consistent with a recent study detailing the role of pseudopaline in zinc uptake (52). Notably, analysis of the *cntO* mRNA by real time PCR (qPCR) demonstrated that the operon is induced under low iron conditions (**Supplementary Materials, Fig S4**). Therefore, the variable impacts of iron on Cnt protein levels observed in the present study may be due to post-transcriptional regulatory mechanisms. These results are consistent with what has been found for the structurally similar staphylopine metallophore produced by *Staphylococcus aureus*, which is regulated both by both iron and zinc (53), suggesting that upregulation of zinc uptake upon iron starvation is a widespread phenomenon.

**Figure 7.**
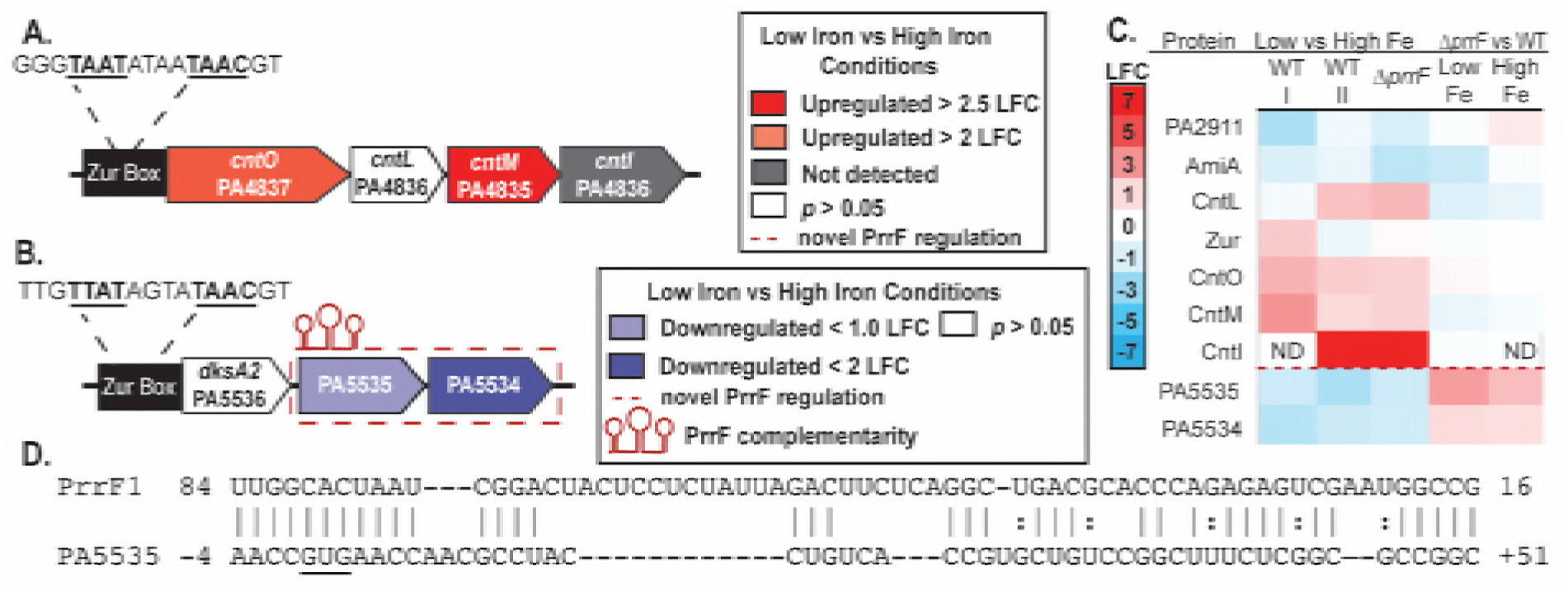
Iron regulates the levels of several zinc-responsive proteins. **A)** Genetic organization of the *cnt* operon. Previous studies identified Zur-dependent zinc regulation of the *cnt* operon, which is hypothesized to occur via a putative Zur box in the *cnt* promoter (54). Both CntO and CntM were found to be upregulated by iron starvation, indicated by orange shading. **B)** DksA1-PA5535-PA5534 genetic organization. The *dksA2* operon is regulated by zinc through the activity of Zur, and a Zur box was identified in the *dksA2* promoter (54, 82). While DksA2 was unaffected by iron, iron activation was shown for the PA5535 and PA5534 proteins (indicated by blue shading). This regulation is dependent upon the *prrF* locus, indicated by a red dashed box. **C)** Heatmap showing iron-dependent log-fold changes (LFC) in zinc-responsive proteins in the first proteomics experiment of wild type PAO1 (I) and the subsequent experiment of wild type PAO1 and ∆*prrF* (II). Novel PrrF regulation was identified for PA5535 and PA5534, separated with a red dashed line. D) Complementarity between PrrF and the PA5535 mRNA was identified using CopraRNA (59).

Our results also showed significant upregulation of the zinc uptake regulator Zur, as well as proteins encoded by several genes found to be upregulated in response to ∆*znuA* induced zinc starvation (54). These include AmiA, encoding an N-acetylmuramoyl-L-alanine amidase (55), and several uncharacterized proteins (PA5534, PA5535, and PA2911) (**Fig. 7C**). The AmiA protein in *E. coli* has a strict requirement for zinc (56). Due to the fact that the *P. aeruginosa* AmiA is downregulated under low iron conditions and upregulated under low zinc conditions, it is possible that AmiA is able to use a different metal in *P. aeruginosa*. While most N-acetylmuranomoyl-L-alanine amidases have a strict zinc requirement, more permissive N-acetylmuranomoyl-L-alanine amidases have been identified (57). Additional zinc-responsive proteins that were induced in iron-depleted conditions included PA5534, PA5535, and PA2911 (**Fig. 7C**). PA5535 is a member of the COG0523 subfamily of the G3E family of P-loop GTPases, which is comprised of metallochaperones and metal-insertases. PA2911 was predicted to be part of an iron ABC permease, but it has been previously shown to be zinc regulated with a putative Zur box. Combined, these data suggest that *P. aeruginosa* responds to iron starvation by modulating the expression of other transition metal homeostasis pathways.

### PrrF mediates iron-regulated changes in multiple iron sparing pathways

The PrrF sRNAs play a significant role in mediating post-transcriptional iron regulation of metabolic processes and virulence in *P. aeruginosa* (23–25, 58). We therefore sought to determine if any of the novel iron regulatory activities uncovered in our proteomics study were dependent upon the PrrF sRNAs. This was achieved by repeating the proteomics experiment with the wild type PAO1 strain and isogenic ∆*prrF* mutant grown in DTSB medium with or without iron supplementation (25). Pathway analysis was performed on all proteins that were significantly (*p <* 0.05) induced or repressed at least two-fold (|LFC| ≥1). Consistent with previous studies (24, 25) and our first proteomics dataset (**Supplementary Materials, Dataset S1**), this analysis demonstrated loss of iron regulation of several iron-dependent pathways in the ∆*prrF* mutant, including oxidative phosphorylation, respiratory electron transport chain, the TCA cycle, and carbohydrate metabolic processes (**Table 2**). Some inconsistencies in iron-regulated pathways between the two experiments were noted, including a lack of iron regulation of the heme acquisition and ketone body synthesis like pathways (compare **Table 1** and **Table 2**). This is likely due to changes in the DTSB media composition between the two experiments, as a result of the variability of TSB batches and differences that can occur during dialysis of the media (see Materials and Methods). However, the results of this later experiment were largely consistent with the first experiment (see the heat maps in **Fig. 1-7**), demonstrating the overall reproducibility of our reported results.

**Table 2.**
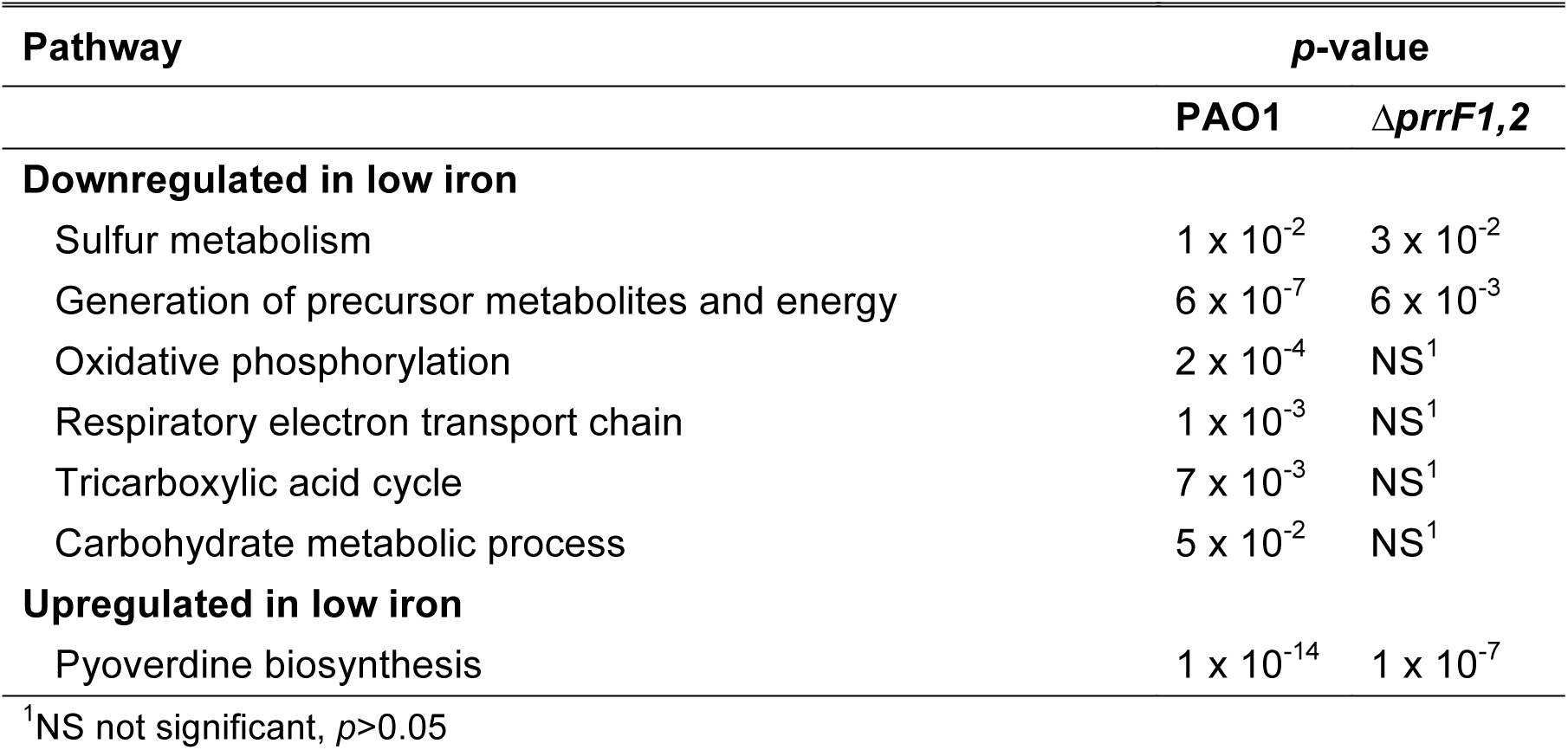
Iron regulated pathways of ∆*prrF1,2* identified through pathway analysis

Notably, this second dataset showed that iron induction of several proteins within newly identified iron-activated pathways was dependent upon the *prrF* locus. Specifically, iron activation of these proteins was lost in the ∆*prrF* mutant, and we observed an increase in the levels of these proteins in the ∆*prrF* mutant compared to wild type when grown under low iron conditions proteins. As expected, PrrF-dependent iron regulation was observed for all previously identified iron-regulated TCA cycle proteins, with the exception of SdhB (**Fig. 1**), as well as for multiple other proteins encoded by PrrF-regulated mRNAs (**Supplementary Materials, Dataset S1**). We additionally found that iron regulation of the branched chain amino acid biosynthesis proteins IlvA1 and IlvD was dependent on the *prrF* locus (**Fig. 2**), as well as several proteins for sulfur metabolism (SsuF, SsuA, CysH, CysN, and CysD; **Fig. 3**) and Fe-S cluster biosynthesis proteins (IscU, and IscS; **Fig. 4**). We further identified novel PrrF regulation of the PhzA phenazine biosynthetic proteins (Fig 5), although it was unclear whether PrrF regulation was occurring through regulation of one or both *phzABCDEFG* transcripts due to the high identity of the encoded proteins. Lastly, we observed PrrF-dependent iron regulation of the zinc-responsive PA5535 and PA5534 proteins (**Fig. 7**). Combined, these results show an even broader impact of PrrF-dependent regulation on iron and metabolite sparing pathways than was previously appreciated, and they highlight a novel role for these sRNAs in mediating metallo-regulatory cross-talk in in *P. aeruginosa*.

### The PrrF sRNAs share complementarity with mRNAs encoding several iron-and PrrF-responsive proteins

To determine whether PrrF is likely to mediate direct regulation of the iron-regulated pathways in this study, we employed CopraRNA (59) to search for complementarity between the PrrF sRNAs and mRNAs encoding PrrF-regulated proteins. Our analysis revealed significant regions of complementarity between the PrrF sRNAs and mRNAs for the TCA cycle enzymes SdhC, PA4333, and PA0794 (**Fig. 1C-E**), none of which were identified as sharing complementarity with the PrrF sRNAs in previous studies. We also identified PrrF complementarity with the mRNA encoding IlvD, which catalyzes two distinct reactions in the branched chain amino acid biosynthesis pathway (**Fig. 2D**). We further identified PrrF complementarity with the mRNA encoding CysD (**Fig. 3C**), encoding the first step in sulfate assimilation into cysteine, and at the 5’ end of *iscS* (**Fig. 4D**). The location of complementarity at the 5’ end of *iscS* is consistent with the discoordinate regulation of the Isc proteins observed in our proteomics analyses (**Fig. 4A,C-D**) and is also consistent with discoordinate regulation of the *isc* regulon in *E. coli* by the iron-responsive RyhB sRNA (7). Lastly, we identified PrrF complementarity with the 5’ UTR and into the coding region of the PA5535 mRNA (**Fig. 7D**), encoding a putative metal chaperone that is induced upon zinc starvation (54, 60) and repressed upon iron starvation (**Fig. 7B-C**). The location of complementarity with the *dksA*-PA5535-PA5534 operon is consistent with the observed discoordinate regulation of these proteins by iron and PrrF, as DksA2 levels were not affected by iron or *prrF* deletion (**Fig. 7B-C**). Combined, these analyses demonstrate the capacity for PrrF to directly interact with mRNAs coding for proteins involved in zinc homeostasis, Fe-S biogenesis, and amino acid metabolism.

### The PrrF sRNAs positively regulate twitching motility

In addition to mediating iron-induced expression of several metabolic and metal homeostasis pathways, the *prrF* locus mediated iron-dependent repression of several twitching motility proteins. Specifically, iron repression of the minor pilins PilE, and FimU, as well as a twitching-associated protein of unknown function PilY2, was eliminated in the ∆*prrF* mutant, and the levels of these proteins were reduced in the ∆*prrF* mutant as compared to wild type grown in low iron conditions (**Fig 6C**). The minor pilin proteins play a key role in initiating twitching motility (61), suggesting the *prrF* locus is required for this activity. In support of this hypothesis, the ∆*prrF* mutant exhibited decreased motility when compared to wild type PAO1 (**Fig. 6B and Supplementary Materials, Fig. S3**). Moreover, iron regulation of twitching motility was abolished in the ∆*prrF* mutant. (**Fig. 6B and Supplementary Materials, Fig. S3**). Thus, our results demonstrate that the *prrF* locus is critical for iron-regulated twitching motility in *P. aeruginosa*, highlighting a novel role for these sRNAs in *P. aeruginosa* physiology and virulence.

## DISCUSSION

*P. aeruginosa* is a versatile opportunistic pathogen that is able to thrive in a variety of nutrient limited environments, including the mammalian host. Despite this organism’s high metabolic iron requirement, *P. aeruginosa* grows proficiently in iron-depleted conditions. This is due in part to the presence of numerous high affinity iron acquisition systems (15), as well as a robust iron-sparing response mediated by the PrrF sRNAs (24, 25). The PrrF sRNAs are already known to contribute to growth in iron limited environments, including the host, as evidenced by multiple studies from our group (23, 58, 62). While the PrrF sRNAs were known to post-transcriptionally repress the expression of non-essential iron-containing proteins to moderate the use of this nutrient, it remained unclear how *P. aeruginosa* could maintain robust growth upon reductions in these pathways. The current study indicates that *P. aeruginosa* uses multiple strategies to compensate for the downregulation of iron containing proteins, including increased production of iron-independent metabolic proteins, metabolite sparing strategies to prioritize production of virulence factors, and increased reliance on zinc as a metal cofactor. Moreover, our study demonstrates that the PrrF sRNAs are responsible for several aspects of these newly-identified responses, providing a mechanistic basis for how *P. aeruginosa* responds to iron starvation. Lastly, our study establishes a novel role for the PrrF sRNAs in iron regulation of twitching motility, providing a potential mechanistic basis for this long-observed phenomenon. As such, this study provides a comprehensive view of how *P. aeruginosa* adapts to iron-limited environments such as the host, and outlines new models for how the PrrF sRNAs contribute to pathogenesis.

One of the overarching themes highlighted by our study is the upregulation of iron-sparing metabolic pathways to compensate for the downregulation of iron-rich metabolic pathways. For example, we show for the first time that iron starvation results in increased production of a putative NADH dehydrogenase, and we confirm the increased production of a cytochrome ubiquinol oxidase (31), which may compensate for PrrF-mediated downregulation of oxidative phosphorylation proteins. The Cyo respiratory pathway uses less iron and can be used without the cytochrome *bc*1 complex (complex III) while retaining the ability to generate a proton motive force by pumping H^+^ into the periplasm (63, 64), and PA2691 encodes a putative type II NADH:quinone oxidoreductase that can regenerate NAD^+^ and maintain redox homeostasis. Together, the Cyo complex and PA2691 have the potential to compensate for the downregulation of the oxidative phosphorylation pathway. The Cyo complex (*bo_3_-*type) is one of five terminal oxidases utilized by *P. aeruginosa*. The others are expressed under starvation (*aa_3_*-type), under high (*cbb_3_*-1 type) or low oxygen (*cbb_3_*-2 type), and under copper limitation (copper independent oxidase, CIO) (63). It was previously shown that *cyoA* expression increases under low iron conditions via loss of Fur repression (31), most likely due to requiring less iron than the other four oxidases. Under iron-replete conditions, the NADH dehydrogenase I, comprised of NuoA-N, is the predominant NADH dehydrogenase as it not only recycles NADH to NAD^+^ but also helps to generate a proton motive force that can be used to generate ATP (39). Likewise, the *cbb_3_*-type cytochrome oxidases are the predominant terminal oxidases, as they interact with the cytochrome *bc*_1_ complex, which also contributes to the proton motive force (63).

Also for the first time, this study shows that *P. aeruginosa* upregulates enzymes in multiple amino acid catabolic pathways when grown in iron-depleted conditions, potentially to compensate for reduced expression of iron-containing TCA cycle enzymes. Pathway analysis identified proteins for ketogenic amino acid metabolism as induced during iron starvation. While ketone body metabolism has not been described in prokaryotes, the induction of branched chain amino acid degradation could indicate a similar metabolic strategy to eukaryotic ketogenesis in response to limited TCA cycle activity. It is likely that the switch to amino acid catabolism is a media-dependent phenomenon, as DTSB is rich in amino acids, which could support this alternative metabolism. This shift is dependent in part on the PrrF sRNAs, which share extensive complementarity with the *ilvD* mRNA and are required for iron-dependent regulation of IlvD and IlvA (**Fig 2D**). A similar shift to amino acid catabolism has been observed in clinical isolates from CF infections as indicated by amino acid auxotrophic mutants (27, 29, 65). These mutants are believed to have lost the ability to synthesize amino acids such as methionine, isoleucine, valine, and leucine, due to the abundance of these amino acids in CF sputum. Host amino acid metabolism is also altered by inflammation (66, 67), potentially contributing the pool of available amino acids in the lung. In this way, inflammation resulting from chronic infection may aid *P. aeruginosa* colonization. Indeed, increased amino acid concentrations are correlated with increased severity of pulmonary disease in CF patients (29). We also found that proteins for amino acid biosynthesis were downregulated upon iron starvation, providing further evidence that *P. aeruginosa* catabolizes amino acids for energy, similar to what is observed during CF lung infections. Thus, iron may play a critical role in the shift of *P. aeruginosa* metabolism toward amino acid catabolism during chronic infection.

In addition to aiding in metabolism, the downregulation of cysteine biosynthesis upon iron depletion may impact the production of Fe-S clusters as cysteine is used as the sulfur donor. Fe-S cluster biosynthesis is mediated by the *iscRSUA-hscBA-fdx2-iscX* operon, which we show here to be discoordinately regulated by iron. Northern blot analysis of the *iscRSUA-hscBA-fdx2-iscX* operon by Romsang, *et al*, showed that these genes in *P. aeruginosa* are transcribed as an operon (68). However, whether these genes are transcribed as an operon under our conditions, or if there is differential regulation of translation by the resulting mRNA transcript, is unknown. In *E. coli*, the *isc* operon is transcribed separately from the *hscBA-fdx2-iscX* operon (41). Thus, it is possible that there is a second promoter resulting in increased expression of HscB, HscA, and IscX independent of the proteins encoded by the upstream genes. The mechanism of the repression of IscS and IscU under low iron conditions is likely attributed to the PrrF sRNAs due to their homology with the *iscS* mRNA translational start site and derepression in ∆*prrF* (**Fig. 4**). To compensate for the downregulation of the desulfonase IscS and the scaffold protein IscU, we identified a partial Suf-like operon encoding a desulfonase, PA3668, and a scaffold protein, PA3667, both of which were upregulated under low iron conditions. In *E. coli* the *isc* operon is considered the housekeeping Fe-S cluster biogenesis operon, but under oxidative stress or low iron conditions the *suf* operon is expressed (41). Our data suggest *P. aeruginosa* uses a similar strategy to maintain limited Fe-S cluster biogenesis under low iron conditions.

The rewiring of metabolic networks that is suggested by our proteomics study also appears to contribute to alterations in virulence factor production. We show that iron starvation downregulates several proteins in the phenazine biosynthesis proteins, and that the PrrF sRNAs negatively affect levels of PhzA, encoded by the first gene in the *phzA-F* operons. No complementarity was identified between the PrrF sRNAs and either of the *phzA* genes or upstream sequence; thus the mechanism for the regulation by PrrF is currently unknown and should be studied more in the future. A possible reason for downregulation of the phenazines under low iron conditions may be to spare chorismate for the production of the siderophore pyoverdine. This metabolite sparing phenomenon was also previously observed with PrrF repression of *antR*, which results in decreased degradation of anthranilate to feed into the TCA cycle, sparing anthranilate for the production of multiple secreted 2-alkyl-4(1*H*)-quinolone metabolites. A further potential example of metabolite sparing for the production of virulence factors is the downregulation of ArgF, which feeds ornithine into the arginine biosynthesis pathway, and the upregulation of PvdA, which incorporates ornithine into pyoverdine. As discussed in the results, ornithine biosynthesis also appears to be upregulated in low iron conditions, possibly to produce enough ornithine for sufficient pyoverdine production.

Many studies have demonstrated that twitching motility increases under low iron conditions, while high iron allows for sessile growth and biofilm formation (50). However, iron-dependent regulation of the genes encoding the twitching motility apparatus has not previously been observed. Here we show that iron starvation increases the levels of several structural components of the twitching apparatus, as well as the major and minor pilin proteins. The fact that these processes have not been identified in previous transcriptional studies could be due to multiple factors, including differences in mRNA and protein half-life or mistmatch between transcription peak and sampling time. Alternatively, these results may indicate a role for post-transcriptional regulatory activities. This latter hypothesis is supported by our subsequent studies of the ∆*prrF* mutant, which lacked the ability to mediate iron regulated twitching motility (**Fig. 6**), highlighting a novel mechanism for how iron may regulate the switch from planktonic to biofilm growth. Future studies will be needed to determine the mechanism by which PrrF promotes the expression of iron-regulated twitching motility proteins.

One notable and somewhat surprising result of our study was iron-dependent regulation of numerous proteins encoded by zinc starvation induced genes, indicating the iron and zinc homeostasis systems in *P. aeruginosa* are integrated. Specifically, the upregulation of proteins in the pseudopaline metallophore system highlights a novel strategy for overcoming iron starvation through the increased expression of zinc acquisition systems. The *cnt* operon is negatively regulated by zinc through the Zur protein via a Zur box in the *cnt* promoter. Thus, it was surprising that this up-regulation occurred when Zur protein levels were also increased under these conditions. One possible explanation for these data is that zinc levels were below the necessary threshold for Zur-dependent repression of the *cnt* operon. In addition to Zur and the pseudopaline proteins, we identified iron regulation of proteins encoded by previously identified zinc-repressed genes: PA5535, PA5534, AmiA, and PA2911 were all induced upon zinc starvation in an earlier study (54), and they were all repressed by iron starvation in our study. We further found evidence for direct discoordinate PrrF regulation of the *dksA2*-PA5535-PA5534 operon, which itself is directly repressed by Zur (54, 70). The interconnection of metal homeostasis systems has not been shown for *P. aeruginosa* but has been shown for other organisms like *Bacillus subtilis* (71). This phenomenon is therefore likely widespread across bacteria and warrants further study in *P. aeruginosa*.

In closing, this study has dramatically increased our understanding of how *P. aeruginosa* regulates metabolism, virulence, and metal homeostasis in response to iron starvation. Previous work has characterized the PrrF dependent iron sparing response, but here we have identified compensatory changes that allow *P. aeruginosa* to continue to thrive when this nutrient is lacking. Further, our results outline several novel roles for the PrrF sRNAs in the iron sparing response, cementing their role as global regulators of metabolism, virulence, and metal homeostasis. With the demonstrated role of iron and the PrrF sRNAs in pathogenesis, these findings will contribute to an increased understanding of how iron regulatory pathways promote *P. aeruginosa* survival in the mammalian host.

## MATERIALS AND METHODS

### Growth conditions

*Pseudomonas aeruginosa* reference strain PAO1 (72) and the isogenic ∆*prrF* mutant (25) were grown in chelex treated dialyzed tryptic soy broth (DTSB) supplemented with 50mM monosodium glutamate and 1% glycerol prepared as previously described (24) supplemented with or without 100 uM FeCl_3_. Cultures were grown at 37°C shaking at 250 rpm. Cells were harvested after 18 hours of growth, the supernatant was removed, and the pellets were stored at -80°C.

### Quantitative label-free proteomics

Cells were lysed in 4% sodium deoxycholate after washing in phosphate-buffered saline. Lysates were washed, reduced, alkylated and trypsinolyzed in filter as previously described (73, 74). Tryptic peptides were separated on a nanoACQUITY UPLC analytical column (BEH130 C18, 1.7 µm, 75 µm x 200 mm, Waters) over a 180 min linear acetonitrile gradient (3 – 43%) with 0.1% formic acid on a Waters nano-ACQUITY UPLC system and analyzed on a coupled Waters Synapt G2S HDMS mass spectrometric system. Spectra were acquired using a data-independent tandem mass spectrometry with traveling wave ion mobility method termed ultradefinition MS^e^ (UDMS^e^). Spectra were acquired using this ion mobility linked parallel mass spectrometry (UDMS^e^) and analyzed as described by Distler *et al.* (30). Peaks were resolved using Apex3D and Peptide3D algorithms (75). Tandem mass spectra were searched against a PAO1 reference proteome (76) and its corresponding decoy sequences using an ion accounting algorithm (77). Resulting hits were validated at a maximum false discovery rate of 0.04. Peptide abundance ratios between the cells cultured in the high iron medium and the cells cultured in low iron medium were measured by comparing the MS1 peak volumes of peptide ions at the low collision energy cycle, whose identities were confirmed by MS2 sequencing at the elevated collision energy cycle as described above. Label-free quantifications were performed using an aligned AMRT (Accurate Mass and Retention Time) cluster quantification algorithm developed by Qi *et al.* (78). Pathways and gene functions were analyzed with information from *Pseudomonas* genome database (76), KEGG database (79) and *P. aeruginosa* metabolome database (PAMDB) (80).

### Twitching motility assays

Twitching motility was quantified as previously described with some modifications (81). Briefly, PAO1 and ∆*prrF* were streaked from freezer stocks onto tryptic soy agar plates and grown overnight at 37°C. Plates of DTSB (prepared as above) with no or 100 µM FeCl_3_, solidified with 1% agar were inoculated using a sterile 10µL pipette tip by stabbing all the way through the agar to the bottom of the plate. The plates were incubated for 18 hours in a humidified 37°C chamber. To visualize the zone of twitching, the agar was removed from the plate and the petri dish was flooded with 1% crystal violet. The crystal violet was incubated for 5 minutes and washed with tap water. The diameter was measured and the average of 10 biological replicates with technical duplicates is presented along with the standard deviation.

### Real-time quantitative PCR

Real-time quantitative PCR (qPCR) analysis of CntO was performed as previously described on five biological replicates of PAO1 grown under the conditions described above. Briefly, relative amounts of cDNA were determined by use of a standard curve generated from serial dilutions of mRNA from PAO1 grown under low iron conditions as described above, which were then reverse transcribed into cDNA. Expression was normalized to *oprF* cDNA in each sample. Primers and probe for *cntO* were as follows: Forward: TTGACAGCGCTCGTATC Reverse: AACTCCGAAGTGGTGAAG Probe: TGTACTCGAACATCGTCAGGCCGC.

## ACKNOWLEDGEMENTS

We thank members of the Oglesby-Sherrouse, Wilks, and Kane laboratories for thoughtful discussion of these studies in laboratory and group meetings. We also thank Dr. Heather Neu and Dr. Sarah Michel for helpful discussion of our results regarding zinc homeostasis.

## LEGENDS FOR SUPPLEMENTARY MATERIALS

**Table S1. Relative protein expression of previously identified iron regulated genes.**

**Figure S1. Detailed view of TCA cycle and aromatic amino acid metabolism showing metabolic intermediates.**

**Figure S2. Detailed leucine degradation (A) and branched chain amino acid degradation pathways (B) showing metabolic intermediates.**

**Figure S3. Representative pictures of twitching motility assay.** The assay was performed using DTSB agar plates supplemented with or without 100 µM FeCl_3_ supplementation solidified with 1% agar. PAO1 twitching motility with iron (**A**) and without iron (**B**). The ∆*prrF* mutant twitching motility with iron (**C**) and without iron (**D**). The scale bar is equal to 10mm.

**Figure S4. CntO expression is upregulated under low iron conditions.** PAO1 was grown for 18 hours in DTSB supplemented with and without 100 µM FeCl_3_, qPCR was performed, and relative expression was determined as described in the Materials and Methods. Expression was normalized to *oprF.* The experiment was performed with *n*=5, *p*=4.94×10^-7^.

**Supplementary Dataset S1. Excel file showing log fold change (LFC) and p values of wild type strain PAO1 grown in low versus high iron conditions.**

